# Targeting host sialic acids in the upper respiratory tract with a broadly-acting neuraminidase to inhibit influenza virus transmission

**DOI:** 10.1101/2023.06.02.543524

**Authors:** Mila B. Ortigoza, Catherina L. Mobini, Hedy L. Rocha, Stacey Bartlett, Cynthia A. Loomis, Jeffrey N. Weiser

## Abstract

The ongoing transmission of influenza A viruses (IAV) for the past century continues to be a burden to humans. IAV binds terminal sialic acids (SA) of sugar molecules present within the upper respiratory tract (URT) in order to successfully infect hosts. The two most common SA structures that are important for IAV infection are those with α2,3- and α2,6-linkages. While mice were once considered to be an unsuitable system for studying IAV transmission due to their lack of α2,6-SA in the trachea, we have successfully demonstrated that IAV transmission in infant mice is remarkably efficient. This finding led us to reevaluate the SA composition of the URT of mice using *in situ* immunofluorescence and examine its *in vivo* contribution to transmission for the first time. We demonstrate that mice express both α2,3- and α2,6-SA in the URT and that the difference in expression between infants and adults contribute to the variable transmission efficiencies observed. Furthermore, selectively blocking α2,3-SA or α2,6-SA within the URT of infant mice using lectins was necessary but insufficient at inhibiting transmission, and simultaneous blockade of both receptors was crucial in achieving the desired inhibitory effect. By employing a broadly-acting neuraminidase (ba-NA) to indiscriminately remove both SA moieties *in vivo*, we effectively suppressed viral shedding and halted the transmission of different strains of influenza viruses. These results emphasize the utility of the infant mouse model for studying IAV transmission, and strongly indicate that broadly targeting host SA is an effective approach that inhibits IAV contagion.

**IMPORTANCE:** Influenza virus transmission studies have historically focused on viral mutations that alter hemagglutinin binding to sialic acid (SA) receptors *in vitro*. However, SA binding preference doesn’t fully account for the complexities of IAV transmission in humans. Our previous findings reveal that viruses that are known to bind α2,6-SA *in vitro* have different transmission kinetics *in vivo*, suggesting that diverse SA interactions may occur during their life-cycle. In this study, we examine the role of host SA on viral replication, shedding, and transmission *in vivo*. We highlight the critical role of SA presence during virus shedding, such that attachment to SA during virion egress is equally important as detachment from SA during virion release. These insights support the potential of broadly-acting neuraminidases as therapeutic agents capable of restraining viral transmission *in vivo*. Our study unveils intricate virus-host interactions during shedding, highlighting the necessity to develop innovative strategies to effectively target transmission.

## INTRODUCTION

In the last century, Influenza A virus (IAV) has led to numerous pandemics, solidifying its enduring presence and burden to humans through continuous transmission. Therefore, there is an urgent need to understand the biology of IAV transmission and devise improved strategies to contain its spread.

Although mice are not the natural hosts for IAV, they have become a valuable tool for studying IAV biology (1–3). However, mouse-to-mouse transmission has historically been inefficient and inconsistent, dampening its appeal as an IAV transmission model. (1, 4–9). Furthermore, mice have been shown to lack α2,6 sialic acid (SA) in their trachea, which are important entry receptors for IAV strains significant to humans.

Avian influenza viruses (AIV) preferentially bind α2,3-SA, which is expressed in the intestine of birds and is a site of viral replication. Recent studies have shown that the tracheal epithelium of birds also express both α2,3-SA and α2,6-SA, indicating a broader distribution of influenza virus receptors in birds than previously known (10, 11). In contrast, humans predominantly express α2,6-SA in the URT, and IAV that preferentially bind α2,6-SA have demonstrated efficient human-to-human transmission. While α2,6-SA is found in parts of the lower respiratory tract (LRT) such as the trachea, bronchi, and bronchioles, it is absent in alveolar cells. In contrast, α2,3-SA is mainly present in the LRT, particularly in alveolar cells, but can also be found in specific cells of the URT (12).

Studies evaluating the composition of SA receptors in the respiratory tract of mice have been limited to LRT-derived cells and tissues. In *ex vivo* murine tracheal epithelial cultures (mTEC), only α2,3-SA was detected (13), while in murine lungs, both α2,3-SA and α2,6-SA were present (14, 15). However, SA distribution in the murine URT (nose, pharynx, larynx), the anatomical compartment where initial infection takes place, has never been evaluated. Furthermore, while there have been indications that α2,6-SA might not be essential for human IAV infection and replication (14), its direct role in transmission was never assessed in an *in vivo* model. Studies of transmission have examined viral mutations of surface viral proteins, receptor binding preferences, and their correlation with transmission in ferrets (16–19). Employing a tractable model like mice to study how host factors contribute to the dynamics of transmission can provide insights into the molecular intricacies of viral-host interactions crucial for our understanding of IAV contagion.

There are key bottlenecks determining transmission in mice which can be categorized into three main stages: virus replication in the URT of the infected (index) host (stage1), release of virus from the URT (shedding) of the infected host (stage2), and arrival of virus in the URT of uninfected (contact) hosts and establishment of productive infection (stage3) (Fig.1). Successfully thwarting transmission requires interruption of any of these stages. Thus, devising vaccines and therapeutics capable of diminishing viral replication, shedding, and/or transmission to contacts is pivotal in our battle against IAV spread.

**Fig. 1.**
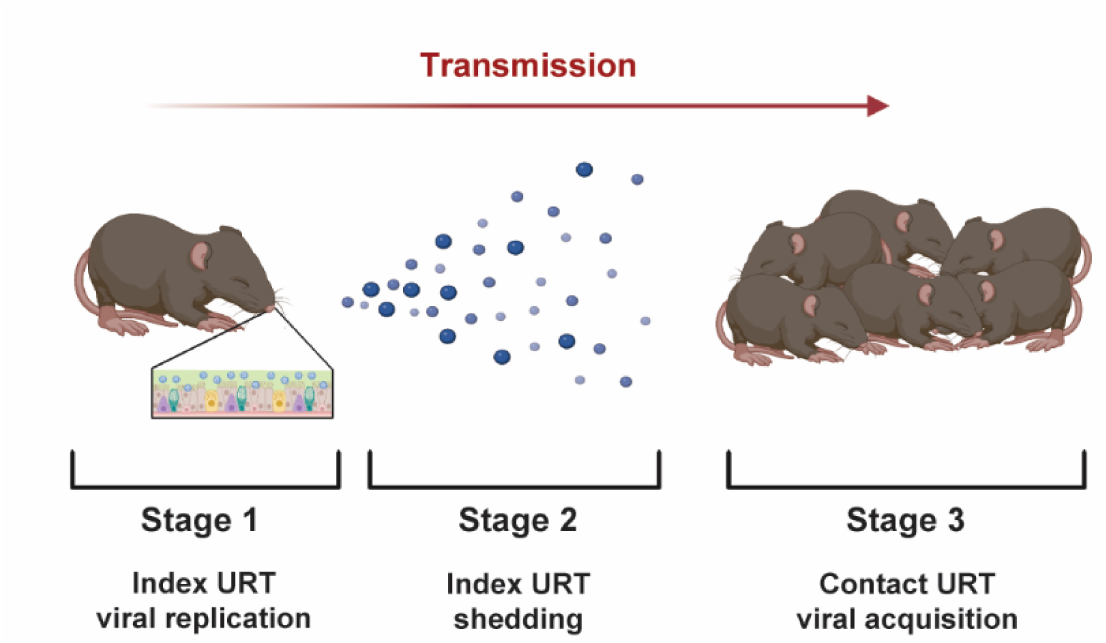
Stages of Transmission. The transmission of influenza virus from index to contact mice involves three distinct stages. First, the virus replicates in the upper respiratory tract (URT) of the infected index mice (**stage 1**). Next, the virus is expelled into the environment (**stage 2**). Finally, virus is acquired by uninfected contact mice and infection is established in the URT (**stage 3**).

Our previous studies illustrate that, in contrast to adult mice, infant mice support efficient transmission of influenza viruses among littermates in an age-dependent and strain-dependent fashion (9). Furthermore, co-infection with *S.pneumoniae* reduced the efficiency of IAV transmission through the activity of bacterial nanA, a broadly-acting neuraminidase, and not nanB, a neuraminidase that selectively targets α2,3-SA. This implied that α2,6-SA may be present and functional in the URT of infant mice. To explore this further, we naturally directed our *in vivo* studies to the URT in order to determine the SA repertoire of both adult and infant mice and evaluate the direct role of SA moieties in IAV shedding and transmission.

## RESULTS

### Mice express both α2,3-SA and α2,6-SA in the URT

Successful transmission of IAV in mice was previously determined to be more efficient in younger mice than older mice (9). This finding exhibited biological plausibility in humans given that children are widely recognized as significant contributors to IAV transmission (20, 21). Hence, we sought to evaluate which stage of transmission was most efficient in infants compared to adults. To achieve this, we infected C57BL/6J mice with A/X-31(H3N2) using a low-volume intranasal (IN) inoculum to prioritize the infection to the URT (7, 22). Infants received 300 plaque-forming units (PFU) in 3μl and adults received 10^5^ PFU in 10μl. After infection, index mice were placed back in the cage with the dam and naïve littermates. Daily shedding samples were collected by gently dipping the nares of each pup in viral medium. Transmission was assessed in contact pups by measuring infectious titers from URT lavages at 4 days post-infection (dpi), a timepoint where >90% transmission is achieved (**Fig.2A**). Interestingly, index URT titers in adults was comparable to that in infants despite being infected with 1000-fold more virus (**Fig.2B**-left). However, the overall shedding titers from days 1-4 was higher in infants compared to adults. This difference was particularly pronounced during the initial two days when peak shedding is expected (**Fig.2B**-middle) (9, 23). Furthermore, acquisition of A/X-31 in contact pups, was significantly higher in infants (92%) compared to adults (33%) (**Fig.2B**-right). Collectively, this suggest that while URT titers were similar between infants and adults at 4dpi, there were notable disparities in shedding (stage2) and in transmission to contacts (stage3). Given that shedding is a correlate of transmission efficiency in this model (9), we focused our studies on virus-host interactions at the epithelium surface. Specifically, we investigated the distribution of SA in the murine URT and their role in IAV shedding and transmission.

**Fig. 2.**
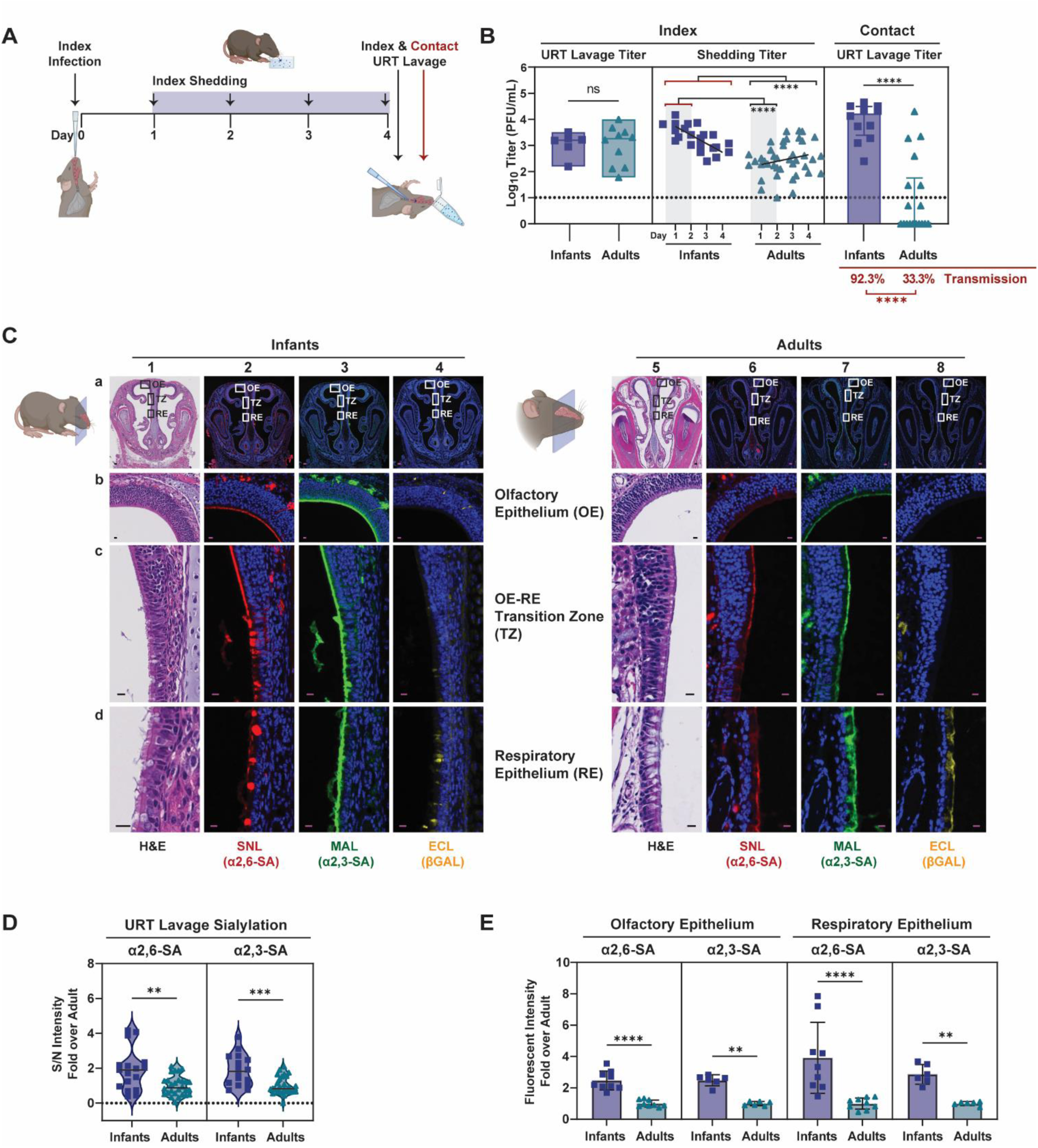
Differential Expression of Sialic Acid in the URT of Mice. **A.** Schematic of Influenza virus transmission. At day 0, index pups were infected IN with A/X-31 (infants at 300 PFU, adults at 10^5^ PFU) and cohoused with uninfected littermates (contact) for 4 days prior to being evaluated for transmission via URT lavage after sacrifice. Shedding was collected on days 1-4 from index pups and infectious virus was quantified via plaque assay. **B.** Transmission of A/X31 in infant and adult mice. Viral titers of lavage fluid and shed secretions were evaluated via plaque assay. Each symbol represents the titer measured from a single mouse, with median values indicated by a line. **C.** Immunohistochemistry of uninfected infants and adult mice nasopharynx. After sacrifice, heads were fixed with 4% paraformaldehyde, paraffin embedded, sectioned through the nasopharynx, and stained with conjugated lectins. Zoomed-in images of the boxed inserts are represented by rows b, c, and d. The scale bars correspond to a length of 100μm in row a, and 10μm in rows b-d. **D.** Uninfected infant and adult mice were subjected to URT lavage, and samples were blotted on a nitrocellulose membrane and immunostained with biotinylated lectins to assess for sialylation of eluted glycoconjugates. Signals were normalized to background and fold over adult mean signal was plotted. Median is denoted by a line. **E.** Fluorescent signal from immunohistochemistry (in C) was measured following color unmixing using ImageJ. Measurements were taken from at least 3 random positions along the epithelial surface of each image, selected from at least 2 representative tissues. Background was subtracted and fold change over the average Adult values were calculated and adjusted for head size. Bars represent median ± interquartile range. **All panels**: Differences among two group medians were analyzed using the Mann-Whitney test. Experiments represent at least 2 biological replicates. Infant mice were 4-7 day old; Adult mice were 8-10 weeks-old; * (p *<* 0.05), ** (p *<* 0.01), *** (p *<* 0.001), **** (p *<* 0.001). Red brackets in statistical comparisons represent the reference group. URT, upper respiratory tract; PFU, plaque-forming unit; H&E, hematoxylin and eosin stain; SNL, *sambucus nigra* lectin; MAL, *maackia amurensis* lectin; ECL, *erythrina crystagalli* lectin; S/N, signal-to-noise ratio; Dotted line denotes limit of detection (10 PFU/mL), where indicated; Zero values were set to 1 PFU/mL.

We first determined whether differences in URT SA levels might account for the variations in shedding and transmission efficiencies between infant and adult mice. URT histological sections from naïve C57BL/6J mice were examined for presence of α2,3-SA and α2,6-SA using conjugated *maackia amurensis* lectin (MAL), *sambucus nigra* lectin (SNL), or *erythrina crystagalli* lectin (ECL), to detect α2,3-SA, α2,6-SA, or exposed β-galactose after desialylation, respectively. *In situ* immunohistochemical (IHC) staining revealed α2,3-SA and α2,6-SA presence in the olfactory and respiratory epithelium of the URT in both infants and adults (**Fig.2C**). The signal for desialylated β-galactose (ECL) was minimal in these naïve animals but prominent during A/X-31 infection (**Fig.4A**). The distribution of α2,3-SA and α2,6-SA in the LRT was also examined (Fig.S1). In both infants and adults, we observed abundant expression of α2,3-SA and α2,6-SA in the lungs, and only α2,3-SA in the trachea, which aligns with previous reports (13–15).

To quantify the levels of α2,3-SA and α2,6-SA in the URT of infants and adults, we performed immunoblots of URT lavages collected after sacrifice. Interestingly, naïve infant mice had higher overall sialylation compared to those of adults by immunoblot of URT lavages and by *in situ* IHC immunofluorescence (IF) quantification (**Fig.2D**-E). Together, these findings indicate that both infant and adult mice possess the necessary terminal SA moieties important for IAV infection, replication, and transmission. Although the expression of sialylated receptors in the URT of adult mice is comparatively lower, viral infection and replication can still take place when a higher virus dose is administered. However, the higher amounts of sialylated receptors in infants appear to contribute, to some extent, to the observed efficiency in index shedding and transmission to contacts in this cohort.

### Blocking both α2,3-SA and α2,6-SA is necessary to inhibit IAV transmission

To evaluate the *in vivo* role of URT α2,3-SA and α2,6-SA in transmission, we infected infant mice with A/X-31 as previously described. Both index and contacts were subsequently treated twice a day (BID) with unconjugated lectins IN for 4 days, with the first treatment starting at ∼6 hours post-infection (hpi). Virus shedding was collected daily and intralitter transmission was assessed in contacts at 4dpi (**Fig.3A**). We observed notable reduction of transmission in all lectin-treated groups in a dose-dependent manner, with simultaneous blockade of both α2,3-SA and α2,6-SA being more effective compared to single SA blockade (**Fig.3B**). Interestingly, selectively blocking α2,6-SA was slightly more effective at inhibiting transmission than α2,3-SA. Overall, these results confirm the presence of functional α2,3-SA and α2,6-SA in the URT of infant mice, underscoring their critical role as host receptors involved in the transmission of A/X-31 in this model.

**Fig. 3.**
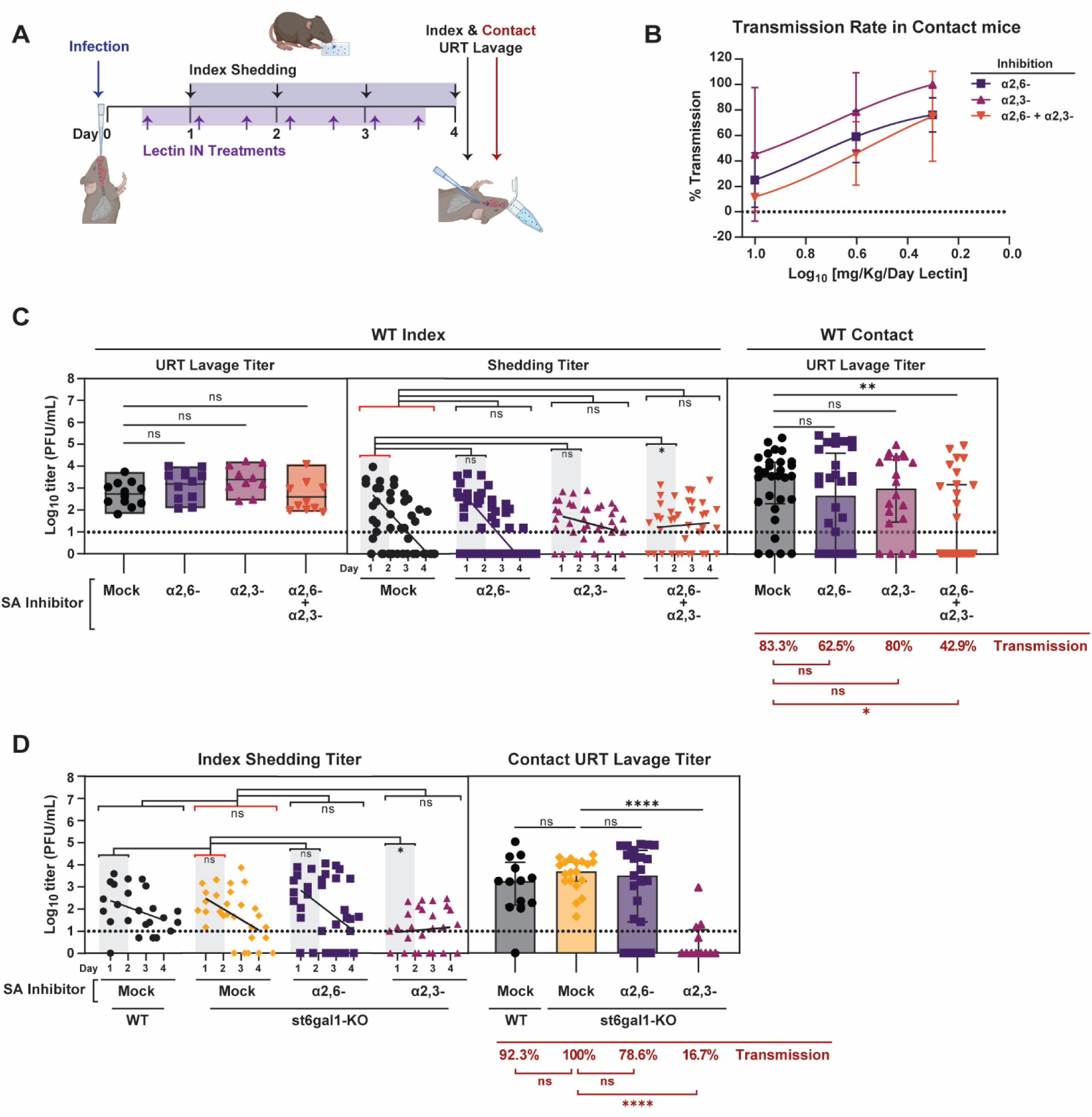
Therapeutic blockade of both α2,3 and α2,6-SA is required to inhibit influenza virus transmission. **A.** Schematic of lectin treatment to block influenza virus transmission. At day 0, pups were infected IN with 300 PFU of A/X-31 (index) and cohoused with uninfected littermates (contact) for 4 days prior to being evaluated for transmission via URT lavage after sacrifice. Shedding was collected on days 1-4 from index pups and infectious virus was quantified via plaque assay. IN lectin treatment was given to both index and contacts, and started 6hpi and continued twice daily until end of experiment. **B.** Transmission rates of at least two independent experiments represented as mean percent transmission with error bars representing the standard deviation. Curves were fit using non-linear regression. **C.** Transmission blockade by lectin inhibitor is represented by virus titers plotted at each stage of transmission. Left panel (stage 1, index URT replication), middle panel (stage 2, index shedding), right panel (stage 3, contact acquisition). **D.** Transmission assay in WT and *st6gal1*-KO infants with or without α2,3-SA inhibitor (MAL) blockade. **Panels C & D**: Median is denoted by a line. within box plots and bar graphs ± interquartile range. Simple linear regression was plotted in shedding panels. Differences among multiple group medians were analyzed using the Kruskal-Wallis test. Experiments represent at least 2 biological replicates. Transmission rates were calculated using Fisher’s Exact test. Infant mice were 4-7 day old; * (p *<* 0.05), ** (p *<* 0.01), *** (p *<* 0.001), **** (p *<* 0.001). Red brackets in statistical comparisons represent the reference group. URT, upper respiratory tract; PFU, plaque-forming unit; IN, intranasal; Mock group represents the vehicle-treated (placebo) group. Dotted line denotes limit of detection (10 PFU/mL), where indicated; Zero values were set to 1 PFU/mL.

We next examined the impact of SA blockade at different stages of transmission. To avoid potential molecular crowding in the URT from higher lectin concentrations, we used a mid-range dose of 4mg/Kg/day. In stage 1, blocking α2,3-SA and/or α2,6-SA did not affect index URT titers (**Fig.3C-left**). In stage 2, the overall shedding titers during days 1-4 were similar between mock-infected infants and those with α2,3-SA and/or α2,6-SA blockade (**Fig.3C-middle**). However, during the initial two days when peak shedding is expected, there was a significant decrease in shedding only when both α2,3-SA and α2,6-SA were simultaneously blocked. In stage 3, selectively blocking either α2,3-SA or α2,6-SA led to a small reduction in URT titers and transmission efficiency in contact mice. However, simultaneously blocking both α2,3-SA and α2,6-SA resulted in the most significant decrease of URT titers and transmission to naïve contacts (**Fig.3C-right**). Collectively, these findings suggest that the presence of URT SA in index mice is essential for effective shedding (stage2), and their presence in contact mice is crucial for efficient virus acquisition (stage3) during exposure.

To investigate the specific role of α2,6-SA in shedding (stage2) and transmission to contacts (stage3), we used mice lacking the gene *st6gal1* (*st6gal1*-KO) which results in the absence of β-galactoside α2,6-sialotransferase 1 (ST6GAL1) (24). This enzyme catalyzes the addition of α2,6-SA to terminal β-galactose of glycoproteins and glycolipids present in mucosal surfaces. Hence, we confirmed the absence of α2,6-SA throughout the URT of *st6gal1*-KO infant mice via IHC (**Fig.S2**) as it has been previously shown for the LRT (14). Interestingly, the shedding and transmission efficiency of A/X-31 in *st6gal1*-KO was comparable to that in wild-type (WT) C57BL/6J mice (**Fig.3D**). To examine if residual α2,6-SA added by a second transferase, the ST6GAL2 enzyme, might be contributing to this phenotype, we administered SNL treatment to both index and contact *st6gal1*-KO to provide additional α2,6-SA blockade. Nevertheless, shedding from index mice remained unaffected, and transmission to contact mice was slightly reduced from 100% to 78.6%. This confirmed that the absence of α2,6-SA alone is insufficient to prevent viral shedding and transmission, as was previously shown for viral replication (14). We next determined if dual SA blockade was necessary to inhibit shedding and transmission by complementing the α2,6-SA deficiency in the *st6gal1*-KO mice with additional α2,3-SA blockade *in trans*. We again found that when both α2,3-SA and α2,6-SA are unavailable in the URT, the index shedding kinetics is altered and inhibited during peak viral shedding on days 1-2, and transmission to contacts is substantially reduced from 100% to 16.7%. These findings again indicate that the presence of both α2,3-SA and α2,6-SA is necessary for efficient shedding and transmission of IAV.

### *In vivo* URT desialylation of both α2,3-SA and α2,6-SA inhibits IAV transmission

DAS181 is a recombinant bacterial NA with enhanced bioavailability *in vivo*. It contains the enzymatic domain from *Actinomyces viscosus* NA fused to the epithelium-anchoring domain of the amphiregulin protein (25). DAS181 is a broadly-acting neuraminidase (ba-NA) promoting desialylation of both α2,3-SA and α2,6-SA and inhibiting influenza viruses (26–29). This is in contrast to zanamivir, a NA inhibitor (NA-I), which targets the viral NA and prevents desialylation of host surface SA. DAS181 has been shown to desialylate murine lungs effectively, yet its effect on the URT had not been evaluated. Hence, naïve infants were either infected with A/X-31 or treated with ba-NA via IN administration for 2 days before sacrifice. URT desialylation was assessed by *in situ* IHC via lectin staining or by URT lavage immunoblots (**Fig.4A-C**). Pups infected with A/X-31 showed detectable signals for IAV nucleoprotein (NP) in the olfactory and respiratory epithelium. As expected, both A/X31-infected or ba-NA-treated pups had increased desialylated β-galactose. Together, these results confirm that ba-NA is as effective as IAV infection in desialylating SA within the URT of pups.

**Fig. 4.**
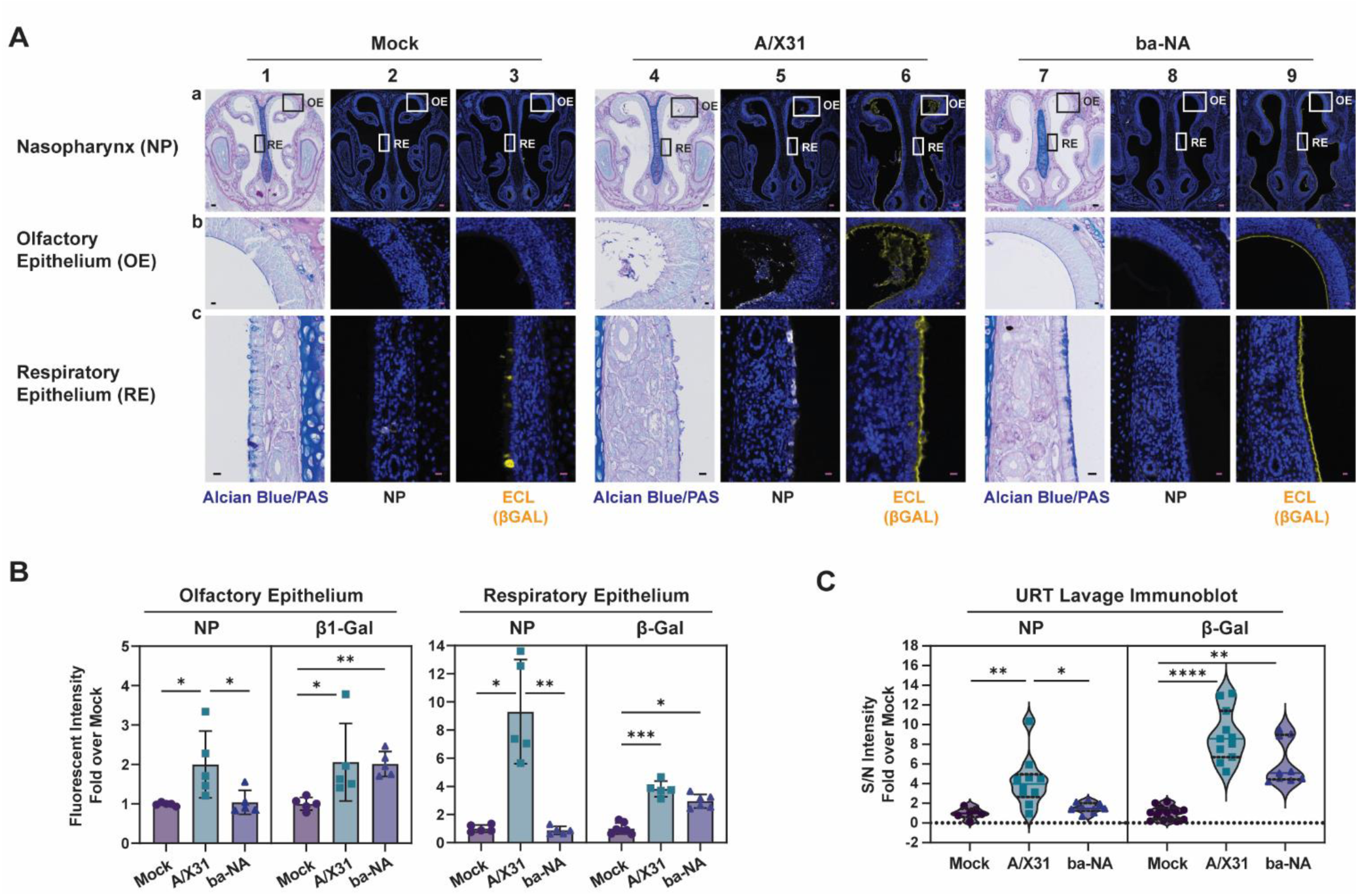
*In vivo* desialylation with broadly-acting neuraminidase. **A.** Immunohistochemistry of uninfected, A/X31-infected, or ba-NA-treated infant mice nasopharynx at 0.5mg/Kg/day. After sacrifice, heads were fixed with 4% paraformaldehyde, paraffin embedded, sectioned through the nasopharynx, and stained with Influenza anti-NP antibody (white) or conjugated ECL (yellow). Zoomed-in images of the boxed inserts are represented by rows b, and c. The scale bars correspond to a length of 100μm in row a, and 10μm in rows b-c. Experiments represent at least 2 biological replicates. Infant mice were 4-7 day old; AB/PAS, alcian blue and periodic acid-Schiff stain; NP, influenza nucleoprotein antibody; ECL, *erythrina crystagalli* lectin. **B.** Fluorescent signal from immunohistochemistry (in A) was measured following color unmixing using ImageJ. Measurements were taken from at least 3 random positions along the epithelial surface of each image, selected from at least 2 representative tissues. Background was subtracted and fold change over the average Mock values were calculated. Bars represent median ± interquartile range. **C.** Mock or ba-NA-treated or A/X31-infected pups were subjected to URT lavage, and samples were blotted on a nitrocellulose membrane and immunostained with biotinylated-Flu-M antibody or biotinylated-ECL to assess for viral infection or desialylated β-galactose in lavage fluid, respectively. Signals were normalized to background and fold over Mock mean signal was plotted. Median is denoted by a line. **All panels**: Differences among two group medians were analyzed using the Mann-Whitney test, and multiple comparisons was done using the Kruskal-Wallis test. Experiments represent at least 2 biological replicates. Infant mice were 4-7 day old; * (p *<* 0.05), ** (p *<* 0.01), *** (p *<* 0.001), ****(p *<* 0.001). URT, upper respiratory tract; Mock group represents the vehicle-treated (placebo) group; ba-NA group represents the DAS181-treated group. AB/PAS, alcian blue and periodic acid-Schiff stain; NP, Flu-NP antibody stained group, β-gal, ECL, *erythrina crystagalli* lectin-treated group; S/N, signal-to-noise ratio; Dotted line denotes limit of detection.

To evaluate the therapeutic potential of *in vivo* desialylation on IAV transmission we first infected infant mice with A/X-31 as previously described, then treated both index and naïve contact pups IN daily with ba-NA or NA-I for 4 days, with the first treatment starting at ∼2hpi. Virus shedding was collected daily and intralitter transmission was assessed at 4dpi (**Fig.5A**). The inhibition of A/X-31 transmission was observed in a dose-dependent manner for both ba-NA and NA-I, and exhibited different inhibitory profiles (**Fig.5B**). To determine the therapeutic window of ba-NA, the inhibition of transmission was evaluated when pups were treated at delayed intervals after infection. Infant mice were infected with A/X-31 then treated with ba-NA or NA-I as previously described. The first cohort received treatment starting at 2hpi, the second at 24hpi, the third at 48hpi, and the fourth at 72hpi. Transmission was assessed in contacts at 4dpi. Transmission inhibition by ba-NA was 100%, 86.7%, 35.8%, and 27.3% when treatment was initiated at 2, 24, 48, and 72hpi, respectively (**Fig.5C**). NA-I exhibited a similar transmission inhibition profile to ba-NA, with rates of 89.7%, 74.3%, 35.9%, and 29.5%, respectively. Next, we analyzed the effects of delayed ba-NA treatment at each transmission stage individually. In stage 1, delayed treatment with ba-NA had no significant impact on URT viral titers (**Fig.5D-left**). In stage 2, viral shedding was reduced only when treatment with ba-NA treatment was initiated early after infection, specifically prior to 24 hours (**Fig.5D-middle**). In stage 3, ba-NA was effective at preventing viral transmission to contacts by >40% if administered within 48 hr of exposure to the infected index (**Fig.5D-right**). Collectively, these findings indicate that broad URT desialylation lowers index shedding and contact transmission in a time-dependent manner, and is most effective within 48hrs of index infection.

**Fig. 5.**
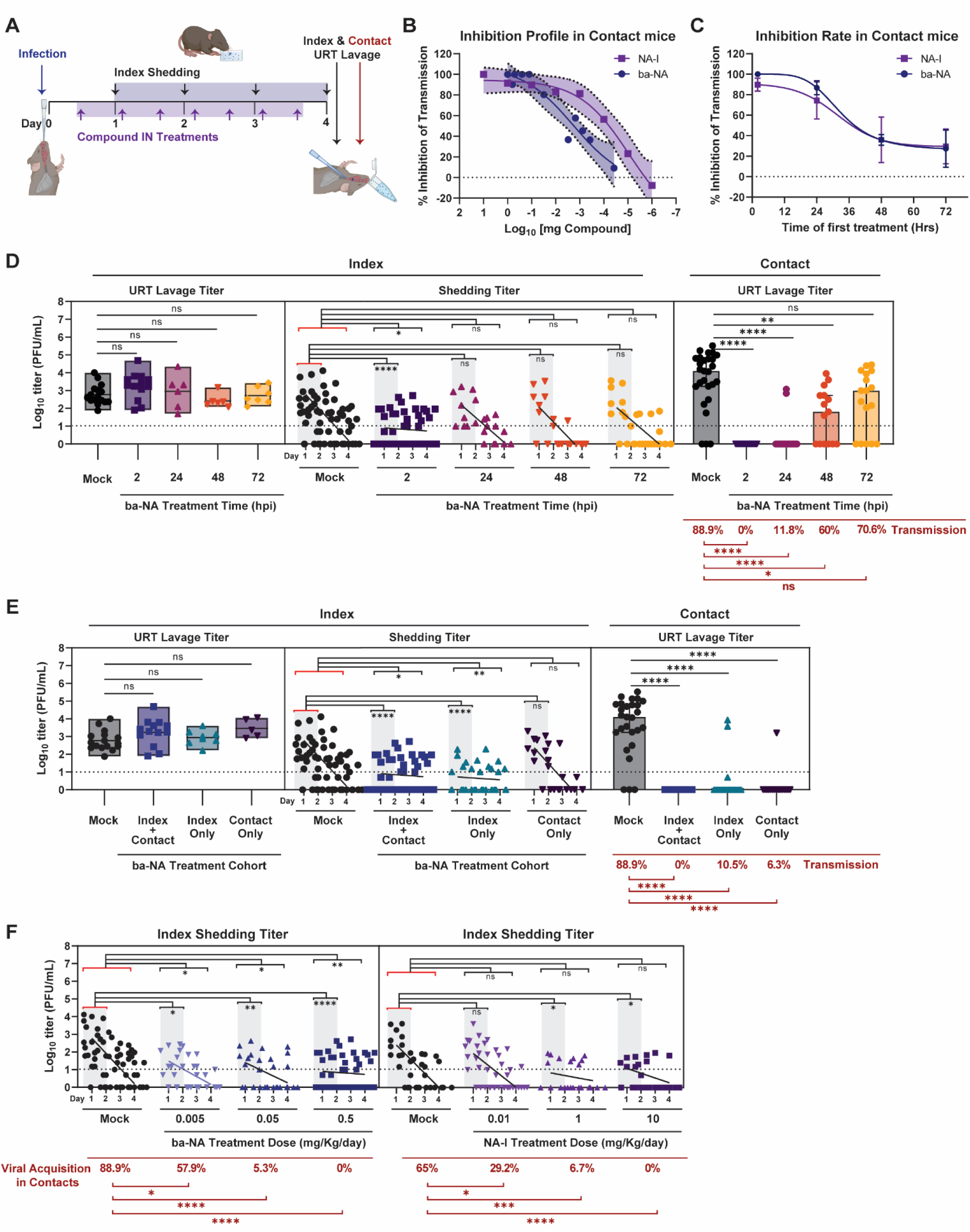
Broadly-acting neuraminidase inhibits influenza virus transmission *in vivo*. **A.** Schematic of compound treatment to block influenza virus transmission. At day 0, pups were infected IN with 300 PFU of A/X-31 (index) and cohoused with uninfected littermates (contact) for 4 days prior to being evaluated for transmission via URT lavage after sacrifice. Shedding was collected on days 1-4 from index pups and infectious virus was quantified via plaque assay. IN compound treatment was started 2hpi and continued once daily for ba-NA (DAS181) or twice daily for NA-I (zanamivir) until end of experiment. **B.** Transmission inhibition rate in contact mice represented as percent (%) inhibition over decreasing compound (ba-NA or NA-I) concentration. % Inhibition was calculated using the following formula: y=100*(1-(y/91.7)) for ba-NA, where 91.7 is the average transmission rate of mock (vehicle-treated) litters, and y=100*(1-(y/65)) for NA-I, where 65 is the average transmission rate of mock (vehicle-treated) litters. Mean transmission is plotted with error bars representing the standard error of the mean (shaded). **C.** Transmission inhibition rate in contact mice represented as percent (%) inhibition over increasing time to compound (inhibitor) treatment. % Inhibition was calculated using the formulas described in (B). Mean transmission is plotted with error bars representing the standard error of the mean. **D.** Transmission inhibition by ba-NA-treatment time is represented by virus titers plotted at each stage of transmission. Left panel (stage 1, index URT replication), middle panel (stage 2, index shedding), right panel (stage 3, contact acquisition). **E.** Transmission inhibition by ba-NA-treatment of individual cohorts is represented by virus titers plotted at each stage of transmission. Left panel (stage 1, index URT replication), middle panel (stage 2, index shedding), right panel (stage 3, contact acquisition). **F.** Shedding titers of A/X31-infected index pups treated with increasing compound (ba-NA or NA-I) concentration. **All panels**: Median is denoted by a line. within box plots and bar graphs ± interquartile range. Simple linear regression was plotted in shedding panels. Differences among multiple group medians were analyzed using the Kruskal-Wallis test. Experiments represent at least 2 biological replicates. Transmission rates were calculated using Fisher’s Exact test. Infant mice were 4-7 day old; * (p *<* 0.05), ** (p *<* 0.01), *** (p *<* 0.001), **** (p *<* 0.001). Red brackets in statistical comparisons represent the reference group. URT, upper respiratory tract; PFU, plaque-forming unit; IN, intranasal; Mock group represents the vehicle-treated (placebo) group. Dotted line denotes limit of detection (10 PFU/mL), where indicated; Zero values were set to 1 PFU/mL.

To determine which cohort had the most influence on IAV transmission, we treated index and/or contacts with ba-NA, and analyzed each transmission stage individually. Compared to mock, URT viral titers in the index was unaffected by any of the treatment combinations (**Fig.5E-left**). Interestingly, overall shedding was reduced only when index pups received ba-NA treatment (**Fig.5E-middle**), consistent with our prior observation that dual receptor blockade using lectins resulted in reduced index shedding (**Fig.3C-middle**). Furthermore, viral acquisition in contact pups was significantly inhibited when either the index, the contact, or both were targeted for URT desialylation, resulting in transmission efficiencies of 10.5%, 6.3%, and 0%, respectively (**Fig.5E-right**). Together, these findings suggest that SA presence in the URT of index pups is important for effective viral shedding, and that inhibiting IAV transmission can be achieved by reducing index shedding (stage2) and/or viral acquisition by contacts (stage3).

While it was anticipated that depleting α2,3-SA and α2,6-SA in the URT of contacts would prevent viral infection upon exposure, the unexpected finding was that depleting SA receptors in the URT of index pups also resulted in decreased viral shedding. Virus elution from the URT surface is not known to necessitate the presence of sialylated glycoconjugates. In contrast, it is known that the cleaving of terminal SA by viral NA promotes virus release (30). To investigate if the presence of SA and its interaction with the virus during egress is an important requirement for efficient shedding from the URT, we targeted host SA using ba-NA at different concentrations and assessed shedding and transmission of A/X31. Increasing the dose of ba-NA resulted in a corresponding decrease in overall shedding dynamics, peak shedding (days 1-2), and viral acquisition in contacts in a dose-dependent manner (**Fig 5F-left**). To confirm that the dissociation of the virus from SA is also important for shedding, we inhibited the ability of viral NA to cleave SA using NA-I at various concentrations and assessed shedding and transmission. As the dose of NA-I increased, the overall shedding dynamics did not show significant changes compared to the mock group, but both peak shedding (days 1-2) and transmission decreased correspondingly (**Fig 5F-right**). Collectively, these findings suggest that the mechanism of viral shedding from the URT is dependent on virion attachment to surface SA during egress, followed by virion detachment from SA during elution.

### *In vivo* URT desialylation does not enhance IAV URT replication or transmission

Ba-NA functions by depleting SA from the surface of cells, thereby preventing important pathogen interaction. However, there exists a possibility that URT desialylation might lead to unintended enhancement of transmission by facilitating the release of progeny virions by complementing viral NA in the catalysis of surface SA (31–33). To address this concern, we first verified that the quantity and size of A/X-31 plaques were diminished in the presence of ba-NA using a plaque reduction assay in MDCK cells (**Fig.6A**). This indicated that ba-NA directly hampers viral replication and spread, aligning with earlier observations from cell-protection assays (25).

**Fig. 6.**
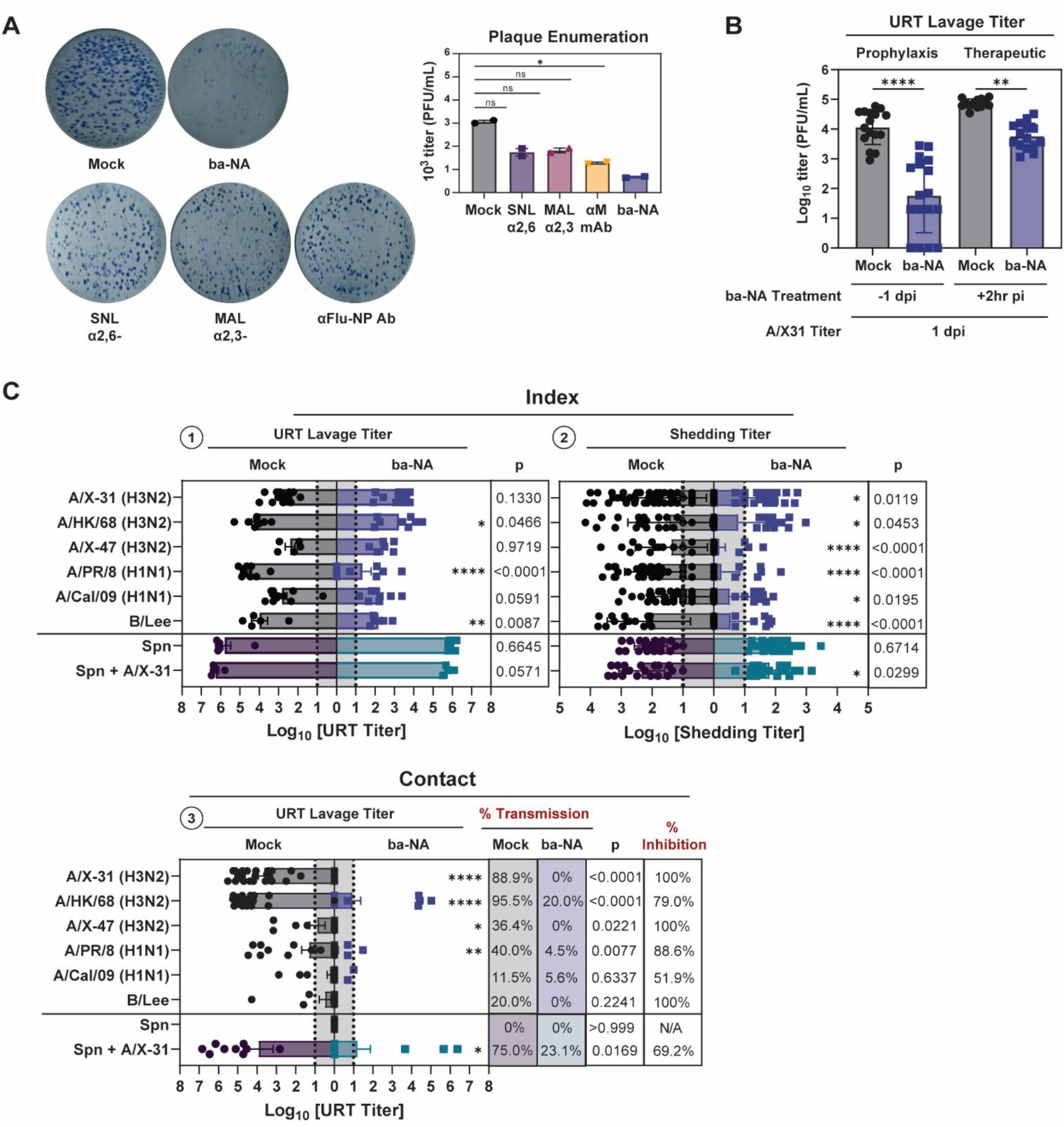
Broadly-acting neuraminidase does not enhance transmission of respiratory pathogens. **A.** Plaque reduction assay testing the inhibitory capacity of ba-NA, lectins, and anti-Flu-NP antibody against A/X31. MDCK cells were infected with equal dose of A/X-31 across each condition, and overlayed with agar containing individual compounds at 2mg/mL for 48-72 hr. Cellular monolayer was fixed with 4% paraformaldehyde and then immunostained with Flu-NP antibody. Plaques were enumerated manually. **B.** Prophylaxis and therapeutic capacity of ba-NA. Pups in a litter were treated with ba-NA 1 day prior or 2 hrs after A/X-31 infection with 300 PFU. Virus titers were assessed at 1dpi by URT lavage after sacrifice. **C.** Transmission of multiple influenza virus strains and Spn in the presence or absence of index-treatment with ba-NA. Index pups in a litter were infected IN with 300 PFUs of selected viruses (or 1000 CFU Spn). Only index pups were treated IN daily with ba-NA for 4 days, with the first treatment starting at 2hpi. Shedding of virus was collected daily and intralitter transmission was assessed in contact littermates by measuring virus from URT lavages at 4dpi. Viral (and Spn) titers were assessed in mock (black for virus, purple for Spn) and ba-NA-treated groups (blue for virus or green for Spn) at the different stages of transmission. **All panels**: Differences among two group medians were analyzed using the Mann-Whitney test, and multiple comparisons was done using the Kruskal-Wallis test. Experiments represent at least 2 biological replicates. Infant mice were 4-7 day old; * (p *<* 0.05), ** (p *<* 0.01), *** (p *<* 0.001), **** (p *<* 0.001). URT, upper respiratory tract; PFU, plaque-forming unit; CFU, colony-forming unit; Mock group represents the vehicle-treated (placebo) group; ba-NA group represents the DAS181-treated group; Spn, *Streptococcus pneumoniae*; MDCK, Madin-Darby canine kidney cells; NP, Flu-NP antibody stained group, Dotted line denotes limit of detection.

Next, we evaluated the preventive effect of ba-NA on primary infection *in vivo*. Pups in a litter were pre-treated IN with PBS or ba-NA 1 day prior to A/X-31 IN challenge with 300PFU. After sacrifice at 1dpi, virus titers were assessed on URT lavage samples. Remarkably, pre-treatment with ba-NA led to a significant inhibition of URT infection following IN challenge, and in some cases, pups were completely protected from infection (**Fig.6B**). An inhibitory effect on URT viral replication in the presence of ba-NA was also observed when the treatment was administered 2hpi and URT replication assessed at 1dpi. In both prophylaxis and therapeutic scenarios, ba-NA treatment did not enhance virus replication in the URT.

We proceeded to investigate whether ba-NA could enhance the transmission of other influenza virus subtypes that inherently exhibit a low transmission (LT) phenotype in this model (9). Index pups in a litter were first infected IN with selected viruses with 300PFU, as previously described. Only index pups then received daily IN treatment with ba-NA for 4 days, starting at 2hpi. Virus shedding was collected daily and intralitter transmission was assessed at 4dpi in mock (black-circles) and ba-NA-treated (blue-squares) groups at the different stages of transmission (**Fig.6C**). The impact of ba-NA treatment on index URT titers (stage 1) varied among viral strains, with some displaying greater susceptibility to ba-NA than others, although none resulted in enhanced URT replication at 4dpi (**Fig.6C-1**). Conversely, index shedding titers (stage 2) were significantly reduced across all viral strains tested upon ba-NA treatment (**Fig.6C-2**). Consistently, titers in untreated contact pups were also lower for all viral strains in the ba-NA index treatment group, particularly for those strains exhibiting moderate and high transmission rates in mock treatment group (**Fig.6C-3**). Notably, there was no enhancement observed in any virus at any transmission stage following ba-NA treatment.

Next, we investigated the impact of ba-NA treatment on the transmission of another respiratory pathogen, *Streptococcus pneumoniae* (Spn). Spn expresses its own NA, enabling it to remove SA from hosts and facilitate immune evasion, attachment, and invasion (34). Furthermore, during co-infection IAV NA has been shown to enhance Spn colonization through the release of free SA which is used by the bacteria for nutrient (35, 36). To assess whether the supplementation of ba-NA could increase URT Spn density, shedding, and transmission to naïve contact littermates, we employed the infant mouse model of Spn transmission (37). Index pups in a litter were first infected IN with 1000 CFU of Spn. Only index pups then received daily IN treatment with ba-NA or PBS (mock) for 4 days. Spn shedding was collected daily by gently tapping the nares of pups onto bacterial plates 10 times. Intralitter bacterial transmission was assessed in contact littermates by measuring Spn levels in URT lavages at 4dpi. Compared to the mock (purple-circles), ba-NA (green-squares) did not enhance URT Spn density, shedding, or transmission (**Fig.6C**). In a separate experiment, Spn-infected index pups were co-infected with A/X-31 at day 3, and bacterial transmission was assessed in contact littermates at 4dpi. Compared to mock, URT Spn density was moderately reduced in the ba-NA treatment group (p=0.0571), which likely resulted in significant Spn shedding reduction (p=0.0299), and Spn transmission efficiency (p=0.0169). These findings suggest that URT desialylation by ba-NA did not result in an increase in Spn transmission at any stage, even if co-infected with IAV. In fact, the effect was inhibitory. Collectively, we demonstrated the efficacy of ba-NA in preventing replication, shedding, and transmission of various IAV strains and Spn in our *in vivo* infant mouse model. Importantly, no enhancement of transmission for any pathogen was observed after therapeutic desialylation in any of the tested conditions in this model.

## Discussion

The interaction of IAV with host SA during viral entry and shedding is complex. The current understanding of IAV transmission efficiency in humans is heavily dependent on viral tropism studies and *in vitro* categorization of viruses as “avian-like” if they preferentially bind α2,3-SA or “human-like” if they preferentially bind α2,6-SA. The SA preference of IAV however, is not sufficient to explain the strain-specific transmission phenotypes observed in humans and in animal models. For example, A/Aichi/68(H3N2), A/Victoria/75(H3N2) all have preferential binding to α2,6-SA in *in vitro* assays (38), yet transmission efficiencies in infant mice is 95% and 36%, respectively (9). Furthermore, the recombinant strain A/X-31(H3N2) which also has preferential binding to α2,6-SA in *in vitro* assays, and was confirmed to contain HA “human-like” residues 226L and 228S (GenBank OQ925911.1), successfully transmits in WT mice and those lacking α2,6-SA (**Fig.3D**) (38–43). This implies that IAV may exploit multiple SA linkages during transmission. Gaining insight into the virus-SA interactions during shedding and entry is essential for our understanding of IAV transmission biology. With the infant mouse model, we can dissect each stage of viral transit and study both viral and host factors important for transmission. In this study, we focused on understanding the contribution of the URT SA repertoire by leveraging mouse biology, and demonstrate the critical role for the host’s sialylated glycoconjugates in shedding and acquisition of influenza viruses *in vivo*.

We confirm that both α2,3-SA and α2,6-SA are present in the URT and lungs of infant and adult mice (**Fig.2C**). Similar to the results in primary mTECs (13), α2,6-SA was not detected in infant or adult mice tracheas (**Fig.S1**). This implies that α2,6-SA expression in the trachea, the conduit to the LRT, is not essential for IAV transmission from the URT. Furthermore, IAV may be more promiscuous in their SA preference than previously thought, given that in the absence of its “preferred” SA, viruses may still utilize “non-preferred” moiety to drive its transmission (**Fig.3C-D**). Our findings illuminate the intricacies of transmission, and underscore the importance of incorporating *in vitro* and *in vivo* assays in future studies to gain a better understanding of the viral-host factors driving IAV transmission.

We also learned that the amount of SA present in the URT of infant and adult mice corresponded with the efficiency of shedding (stage2) and transmission to contacts (stage3) (**Fig.2**). Using two independent methods, we found that adult mice in general had lower amounts of both α2,3-SA and α2,6-SA in the URT compared to infants, which can partially explain the lack of efficient transmission in this cohort. This implies that a threshold of SA molecules is necessary to facilitate effective shedding and ensure sustained infection in the receiving contact. We tested this hypothesis by decreasing the available SA in the URT and demonstrating inhibition of both index shedding and contact transmission using three different and independent approaches: 1) simultaneously blocking both α2,3-SA and α2,6-SA using SA-specific lectins (**Fig.3C**); 2) using an *st6gal1*-KO mouse, which fails to conjugate terminal α2,6-SA in all tissues, complemented with selective α2,3-SA blockade (**Fig.3D**); and 3) simultaneously removing both α2,3-SA and α2,6-SA in the URT using exogenous ba-NA (**Fig.4-5**). Selectively inhibiting either α2,3-SA or α2,6-SA was insufficient at hindering IAV transmission. Hence, broad URT desialylation early during infection and exposure is a promising strategy to decrease the efficiency of IAV transmission.

The observation that decreasing the availability of SA in the URT led to a suppression of virus shedding was surprising because the presence of SA is not traditionally considered essential for virus URT release. In contrast, tissue culture studies have demonstrated that removing SA with exogenous NA promoted viral replication and elution from cells (32). To delve deeper into this discrepancy, we examined the shedding efficiency of IAV under two conditions: one where SA was absent (achieved with ba-NA) and another where the virus-SA interaction was disrupted (achieved with NA-I). Our results indicated that shedding was reduced in a dose-dependent manner in both conditions, suggesting that the production of progeny virions rely on a sequential process involving virus binding to SA to assist virion egress and formation, followed by its release from SA for final elution. This implies that SA could be a prospective host target for future antivirals.

Because of the unexpected success of ba-NA in suppressing shedding and inhibiting transmission of various IAV subtypes (**Fig.6C**), we became interested in its effect on other URT pathogens that are known to cause secondary infections following IAV infection. Considering that IAV-Spn co-infection enhances bacterial colonization and pathogenesis through NA activity, we were apprehensive that ba-NA therapeutics may have inadvertent impacts on co-colonizing bacterial pathogens (36). To investigate this possibility, we evaluated the effects of ba-NA on Spn replication, shedding, and transmission in the presence or absence of co-infection with IAV. Our results indicated that ba-NA significantly reduced Spn shedding in the IAV-infected cohort. Moreover, none of the Spn cohorts, regardless of co-infection status with IAV, exhibited any enhancement of Spn replication, shedding, or transmission to contacts. These results expand upon earlier findings that demonstrated how ba-NA treatment of IAV-infected mice indirectly lowered secondary Spn pneumonia (44). This finding strengthens our confidence in the safety of employing ba-NA as a therapeutic targeting the host to impede IAV shedding and transmission. Furthermore, a host-targeting therapeutic might have a lower risk of developing resistant variant strains compared to viral-targeting compounds. As a result, ba-NA could potentially become the first antiviral agent capable of obstructing viral shedding when directly applied to the nasal mucosa therapeutically, as well as providing protection against viral infection when used prophylactically. Ba-NA inhibitors hold the potential to be a significant advancement in antiviral therapy, offering the possibility of outwitting viral evolution.

## Materials and Methods

### Mice

C57BL/6J (000664, Jackson-Laboratories), and *st6gal1*-KO (Joseph Lau, Roswell Park Cancer Institute, NY) (24) were bred in a conventional animal facility. Pups were housed with their dam for the duration of experiments. Studies and procedures were conducted in accordance with the Guide for the Care and Use of Laboratory Animals (73) and the Biosafety in Microbiological and Biomedical Laboratories recommendations (45), and were approved by the Institutional Animal Care and Use Committee of NYU Langone Health (Assurance A3317-01). KAPA2G HotStart Mix (07961316001, Roche) was used in genotyping.

Fwd-CTGAATGGTGGACTGTGG

*st6gal1*-KO_Rev-CATTTTGTGAGCCCCATTAG

WT_Rev-TGTTGAAGGGAGAATCTGTG. (24).

### Biological Reagents

#### Cells

MDCK cells (CCL-34, ATCC) were cultured in DMEM (HyClone), 10% fetal bovine serum (Peak), and 1% penicillin-streptomycin (Gibco) with monthly mycoplasma testing.

#### Viruses

A/X-31(H3N2) (GenBank:OQ925911-18), A/X-47(H3N2) (NR-3663, BEI), A/Hong Kong/1/1968-2_MA21-2(H3N2) (NR-28634, BEI), A/Puerto Rico/8/1934(H1N1) (NR-3169, BEI), A/California/4/2009(H1N1) (NR-13659, BEI), B/Lee/1940 (NR-3178, BEI). IAV and IBV were propagated in 8-10 day-old embryonated chicken eggs (Charles-River, CT) for 2 days, 37°C and 33°C, respectively. Allantoic fluid was ultra-centrifuged (10,000RPM, 30min, 4°C) and stored at -80°C. Virus titers were quantified via standard plaque assay in MDCK cells in the presence of TPCK-trypsin (20233; Thermo-Scientific) (46).

#### Bacteria

Streptomycin-resistant TIGR4 strain (P2406, provided by J.N.W.) was cultured on tryptic-soy-agar (TSA)+streptomycin (200 μg/mL) (37).

#### Compounds

Broadly-acting neuraminidase (DAS181, Ansun BioPharma, Inc, CA); Zanamivir (SML0492, Sigma); unconjugated-SNL (L-1300-5, Vector-Laboratories); biotinylated-SNL (B-1305-2, Vector Laboratories); SNL-Cy5 (CL-1305-1, Vector-Laboratories); unconjugated-MAL-I (L-1310-5, Vector-Laboratories); biotinylated-MAL-I (B1315-2, Vector-Laboratories); biotinylated-ECL (B-1145-5, Vector-Laboratories); fluorescein-streptavidin (SA-5001-1, Vector-Laboratories).

#### Antibodies

anti-Flu-NP (ab20343, Abcam), biotinylated-anti-Flu (ab20351, Abcam); HRP-anti-mouse-IgG (ab205719, Abcam), streptavidin-HRP (21130; Pierce).

### Infection, shedding, and transmission

Infants (3-7 days old) and adults (8 weeks old) were infected IN with 3μl (300 PFU) or 10μl (10^5^ PFU), respectively, without general anesthesia. Virus shedding was collected by dipping mouse nares in medium (PBS, 0.3% bovine-serum-albumin) daily. Index:contact ratio was 1:2-1:3. Transmission was determined in contacts at 4dpi. Euthanasia via CO_2_ asphyxiation and cardiac puncture preceded URT lavage (300μl PBS infusion through the trachea and collecting via the nares).

### Compound treatments

All compounds were dosed by weight (mg/Kg/day). Lectins were diluted in buffer [10mM HEPES, 0.15M NaCl, 0.1mM CaCl2, 0.08% NaN3], dosed twice-daily; ba-NA (DAS181) was diluted in PBS, dosed daily. NA-I [zanamivir, Sigma] was diluted in water, dosed daily.

### Plaque reduction assay

MDCK cell monolayers were infected with A/X-31 (200 PFU/mL/well) in 6-well-plate. At 45-60 minutes, virus was aspirated and plaque media-containing lectins, DAS181, or anti-Flu-NP (2mg/mL) was overlaid onto monolayer and incubated (37°C, 48-72hrs). Plates were fixed with 4% paraformaldehyde and plaques immunostained with anti-Flu-NP (1hr, 1:1000). Three washes [PBS, 0.5% milk-powder] were done prior to HRP-anti-mouse-IgG (1hr, 1:5000). Three washes were done prior to applying TrueBlue-HRP substrate (5067428, Fisher) and enumeration.

### URT lavages Immunoblots

URT lavages (150μL, 1:50 in TBS [tris-buffered saline]) were applied to slot blotting apparatus (Minifold-II, Schleicher-Schuell) containing pre-wetted 0.2μm nitrocellulose membrane (GE10600094; Amersham). After applying vacuum, membrane was blocked with Carbofree buffer (SP-5040-125, Vector-Laboratories) (1h, RT) followed by incubation with biotinylated-lectins (2μg/mL) or biotinylated-anti-Flu primary (1:10,000) in TBS-T (TBS, 0.1% Tween-20) overnight (4°C). Membrane was washed 5x/10min with TBS-T prior to streptavidin-HRP secondary (1:100,000, 1hr, RT). After 5 washes/10 min with TBS-T, membrane was developed using SuperSignal West-Femto substrate (34095; Thermo-Scientific) using iBright-Imaging (Invitrogen). Mean-gray-values were measured with ImageJ (v.1.53e, NIH), background was subtracted, and normalized to a positive control (URT lavage from A/X-31-infected mice) included on all blots. Fold-change over the average Adult values were plotted.

### Immunohistochemistry

Pups (uninfected, A/X-31-infected, or ba-NA-treated) were euthanized after 2dpi (∼5-6 days old). After skin removal, intact heads were fixed in 4% paraformaldehyde (48-72 h, 4°C). Heads were washed in PBS (4°C, 30 min), decalcified (0.12M EDTA [infants], 0.25M [adults]) (4°C, gentle-shaking, 7-days [infants], 21-days [adults]), dehydrated through graded ethanols, and paraffin-embedded (Leica Peloris). 5μm sections were immunostained (Leica BondRX). Briefly, deparaffinized sections were incubated with Rodent Block (RBM961L, Biocare), then with primary reagents (2hr, Cy5-SNL [1:50], biotinylated-MAL-I [1:100], biotinylated-MAL-2 [1:50], biotinylated-ECL [1:200], or anti-Flu-NP (1hr, [1:500]). Lectins were followed by Streptavidin-fluorescein (1:50), whereas anti-Flu was followed by Mouse-on-Mouse-HRP-conjugated polymer (MM620L, Biocare) and Opal620 (FP1495001T, Akoya-Biosciences). Slides were counterstained with DAPI and scanned (Akoya Polaris Vectra). Images were prepared with OMERO.figure v4.4 (OME team). IF was measured using ImageJ (v.1.53e) following color unmixing. Mean-gray-value were obtained from 3-5 random positions along the epithelial surface of each image of >2 representative tissues. Background fluorescence was measured using the same selection and subtracted from the signal. Fold-change over the average Adult value were calculated and adjusted for head size.

### Statistical analysis

GraphPad Prism (v9.4, CA) was used for all analyses (nonparametric, two-tailed tests at α=0.05).

## Data Availability

Data is available upon request.

## Acknowledgments

We thank Joseph T. Y. Lau for providing *st6gal1*-KO mice, Ansun BioPharma, Inc (Trevor Gale and Stephen Hawley) for providing DAS181 (Fludase^®^), and the NYU Experimental Pathology Laboratory for their collaboration (Mark Alu, Branka Brukner Dabovic, Shanmugapriya Selvaraj, and Valeria Mezzano).

## Author contributions

Conceptualization: M.B.O., Data curation: M.B.O., C.L.M., H.L.R., S.B., Formal analysis: M.B.O., Funding acquisition: M.B.O., J.N.W., Investigation: M.B.O., C.L.M., H.L.R., S.B., Methodology: M.B.O., C.A.L., Project administration: M.B.O., Supervision: M.B.O., Validation: M.B.O., Visualization: M.B.O., Writing-original draft: M.B.O., Writing-review & editing: M.B.O., C.L.M., H.L.R., S.B., C.A.L., J.N.W.

