## Supplemental Figures for "Targeting host sialic acids in the upper respiratory tract with a broadly-acting neuraminidase to inhibit influenza virus transmission"

### Supplemental Figures and Legends

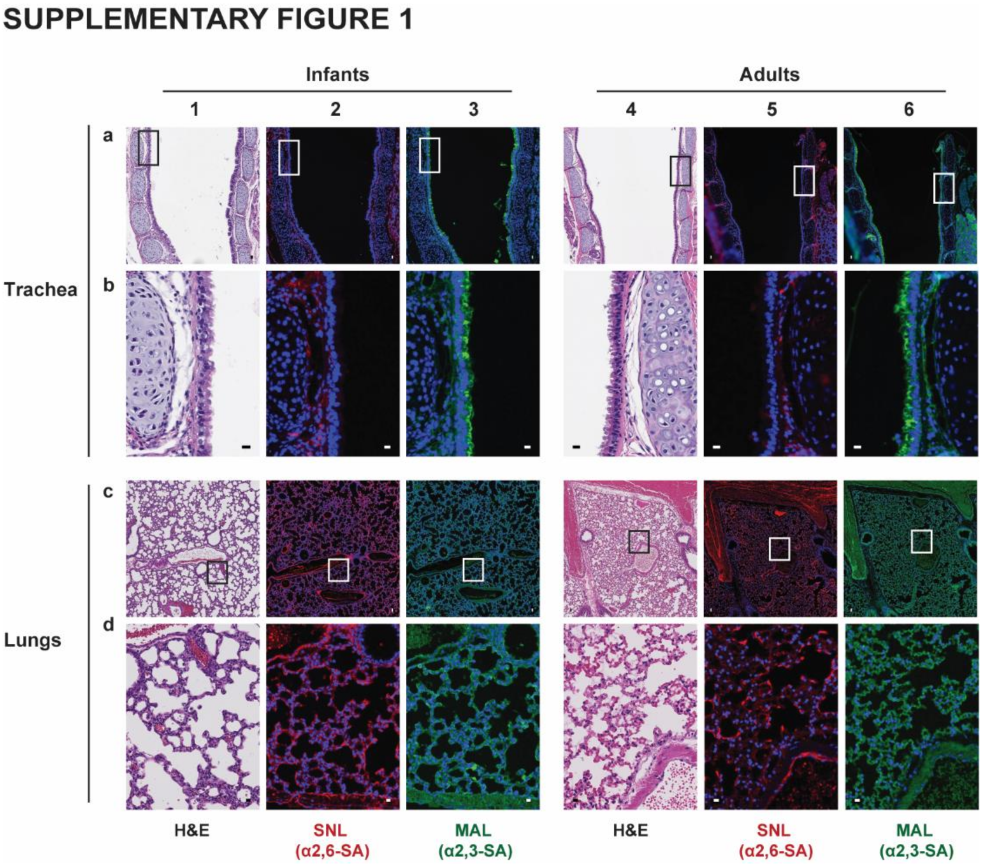


**Fig.S1. Differential Expression of Sialic Acid in the LRT of Mice.**

Immunohistochemistry of uninfected infants and adult mice trachea and lungs. After sacrifice, trachea and lungs were individually dissected, fixed with 4% paraformaldehyde, paraffin embedded, sectioned, and stained with conjugated lectins. Sections displayed represent at least 2 biological replicates. Infant mice were 4-7 day old; Adult mice were 8-10 week-old; LRT, lower respiratory tract; H&E, hematoxylin and eosin stain; SNL, *sambucus nigra* lectin; MAL, *maackia amurensis* lectin; ECL, *erythrina crystagalli* lectin; Zoomed-in images of the boxed inserts are represented by rows b and d. The scale bars correspond to a length of 100μm in rows a and c, and 10μm in rows b and d.

### SUPPLEMENTARY FIGURE 2

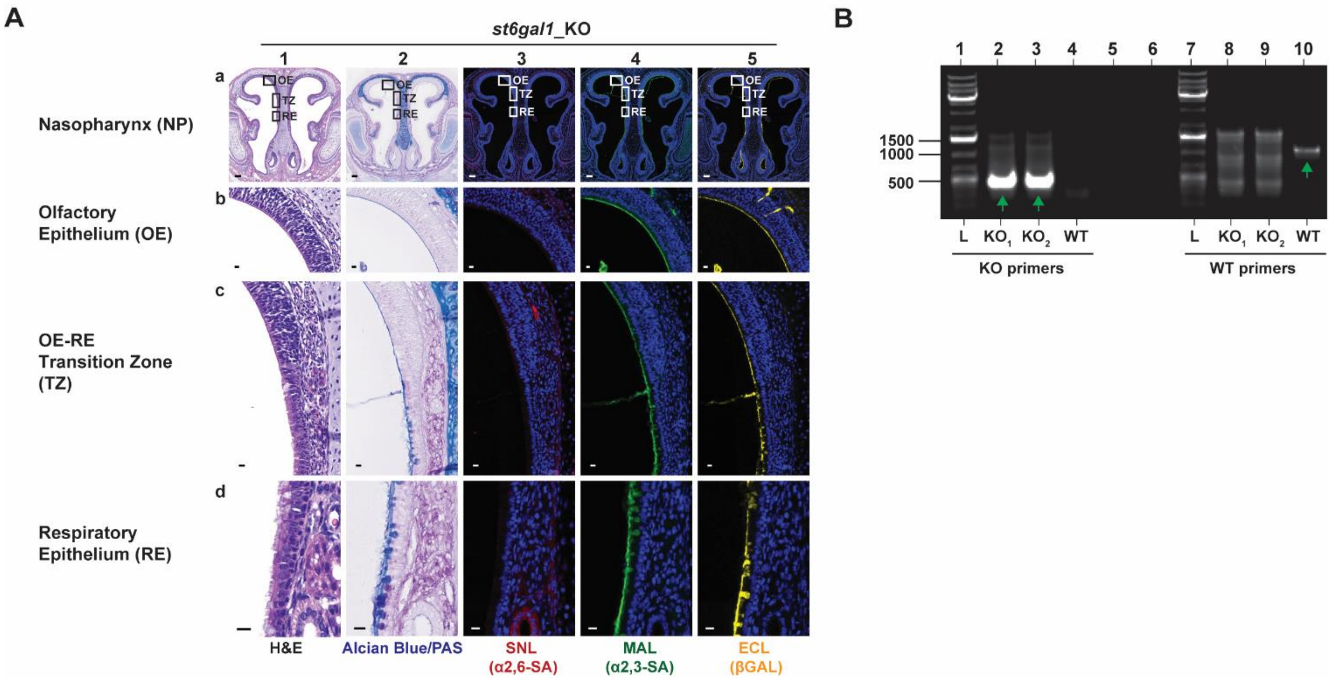


**Fig.S2. *st6gal1*-KO Mice.**

**A.** Immunohistochemistry of uninfected *st6gal1*-KO infant nasopharynx. After sacrifice, heads were fixed with 4% paraformaldehyde, paraffin embedded, sectioned through the nasopharynx, and stained with conjugated lectins. Zoomed-in images of the boxed inserts are represented by rows b, c, and d. The scale bars correspond to a length of 100μm in row a, and 10μm in rows b-d. Experiments represent at least 2 biological replicates. Infant mice were 4-7 day old; H&E, hematoxylin and eosin stain; AB/PAS, alcian blue and periodic acid-Schiff stain; SNL, *sambucus nigra* lectin; MAL, *maackia amurensis* lectin; ECL, *erythrina crystagalli* lectin. **B.** Genotyping of *st6gal1*-KO dam and sire. Ear notches were subjected to DNA isolation, and PCR was done using WT and KO primers specific for the ST6GAL1 gene (1.2 Kb segment). Lack of ST6GAL1 gene would yield a 500 bp segment. Green arrows denote correct bands. Lanes 1 and 7 is the 1Kb plus DNA ladder.
